# Psilocybin alters visual contextual computations

**DOI:** 10.1101/2025.02.06.636848

**Authors:** Marco Aqil, Gilles de Hollander, Nina Vreugdenhil, Tomas Knapen, Serge O. Dumoulin

## Abstract

Psilocybin alters perception and brain dynamics. Contextual computations are ubiquitous in the brain. Here, we investigate the effects of psilocybin using psychophysics, ultra-high field functional MRI, and computational modeling. We find that 1) psilocybin alters contextual perception in the Ebbinghaus illusion, 2) psilocybin alters contextual modulation in cortical responses to visual stimuli, and 3) we propose a computational model capable of capturing and linking these changes. Leveraging vision as a beachhead, our findings highlight the alteration of contextual computations as a potential general mechanism underlying psychedelic action.

**Teaser:** Psilocybin alters visual-contextual computations, a potential general computational mechanism for psychedelic effects in the human brain.

---

Psilocybin is a serotonergic hallucinogen or classic psychedelic. Previous studies have shown that psilocybin alters perception and brain dynamics ^1–5^. Yet, the underlying computational changes remain unclear. Contextual computations are ubiquitous in the brain, and computations first discovered in vision have later been observed in other sensory and cognitive domains ^6–9^. Here, we investigated the effects of psilocybin in 5mg and 10mg doses with psychophysics, ultra-high-field fMRI, and computational modeling in a randomized, double-blind, placebo-controlled, crossover design. We explicitly tested whether and how psilocybin alters visual-contextual computations in brain and behavior, and provided an explicit mathematical model of cortical responses at the single-timecourse level. Our findings highlight the alteration of contextual computations as a potential general and parsimonious computational mechanism underlying the effects of psychedelics on brain and behavior.

## Psilocybin alters contextual perception

To examine the effects of psilocybin on contextual perception, we used the Ebbinghaus illusion ^10,11^. In this classic perceptual illusion, the perceived size of a target stimulus is altered by the presence of a visual context. Previous studies have related the Ebbinghaus illusion to anatomical and functional properties of primary visual cortex (V1) ^11–13^. Here, participants were asked to report which of two concurrently presented visual stimuli was larger, while maintaining fixation on a central cross, following placebo, 5mg, and 10mg psilocybin administration (Fig 1). In control trials, both stimuli were presented in isolation (Fig 1a, bottom-right inset). On test trials, one of the two stimuli was presented within a context of larger stimuli, making it appear perceptually smaller (Fig 1a, top-left inset). Psilocybin significantly increased the Ebbinghaus illusion by 39% (5mg), and 59% (10mg) relative to placebo (Fig. 1b). Relative to true stimulus size, the Ebbinghaus illusion was 6% with placebo, 8.3% with 5mg psilocybin (*p*: 0.012), and 9.6% with 10mg psilocybin (*p*: 0.005) (Fig. 1c). The change in the Ebbinghaus illusion induced by psilocybin could not be explained by potential concurrent changes in noise or lapses, which were also included in the psychophysical model (Methods, Extended Data 1). Perception in control trials (i.e. stimuli presented without context) was unaltered by psilocybin (Fig 1a shaded gray and Extended Data 1), showing that the effect of psilocybin was specific to contextual perception. In sum, we found that psilocybin altered contextual perception, specifically increasing the Ebbinghaus illusion. This finding highlights the alteration of contextual computations as a potential mechanism underlying the effects of psilocybin on perception, and demonstrates a novel neuromodulatory contribution to the Ebbinghaus illusion.

**Fig. 1.**
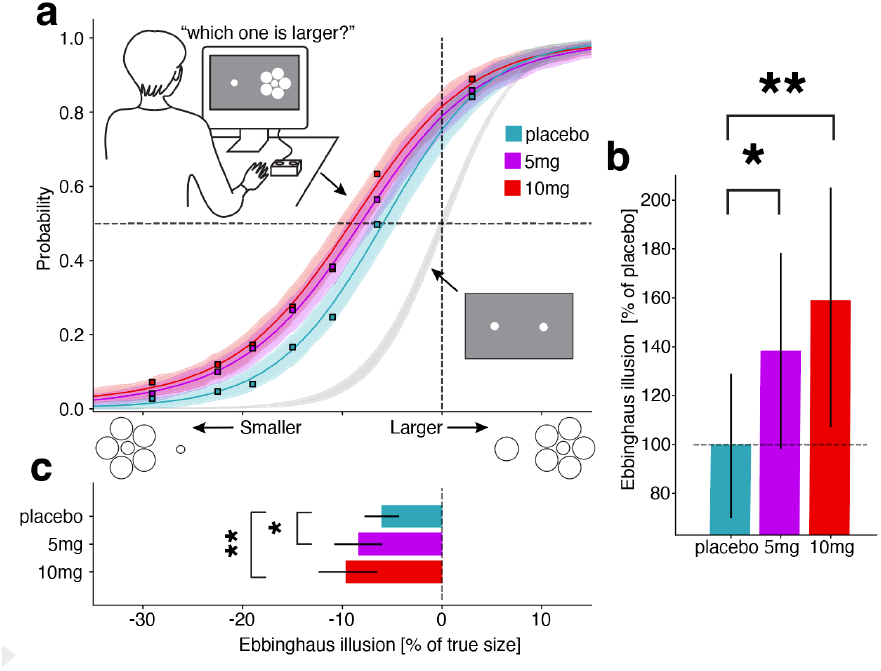
Psilocybin alters contextual perception in the Ebbinghaus illusion. **a**, Participants were asked to judge which one of two concurrently presented stimuli was larger. In control trials, stimuli were presented in isolation(bottom right inset), and perception was unaltered by psilocybin (shaded gray, Extended Data 1). In test trials, one stimulus was presented within a context of larger stimuli (top-left inset), resulting in a smaller perceived size (Ebbinghaus illusion). Psychometric data for test trials (squares), estimated psychometric curves (colored lines), and 95% highest density intervals for the posterior model predictions (shaded color) are shown for placebo (cyan), 5mg (magenta), and 10mg (red) psilocybin doses. **b**, Psilocybin significantly increased the Ebbinghaus illusion by 38% (5mg dose) and 59% (10mg dose) relative to placebo. **c**, Relative to true stimulus size, the Ebbinghaus illusion was 6% with placebo administration, 8.3% with 5mg psilocybin (*p*: 0.012), and 9.6% with 10mg psilocybin (*p*: 0.005). One asterisk indicates *p*<0.05, two *p*<0.01 (Bayesian p-value, one-sided test).

## Psilocybin alters contextual brain responses

To probe the effects of psilocybin on contextual brain responses, we measured 7T fMRI responses to a simple visual stimulus (Fig 2a), a contrast-defined checkerboard bar sweeping across the central 10 degrees of visual field in eight directions (Fig 2b) ^14,15^. Participants maintained fixation, performed a dot-color change task at fixation, and eye movements were recorded (Fig 2b, Extended Data 2). Responses were mapped to individual cortical surfaces, and visual field maps drawn on the basis of individual polar angle and eccentricity maps (Fig 2c) ^16^. Previous studies have identified a variety of contextual modulations such as surround suppression and nonlinear spatial summation in brain responses to simple visual stimuli ^9,15,17,18^. Surround suppression(Fig 2d) is a particular form of contextual modulation, measurable with imaging or recording methods, thought to arise via a combination of lateral and feedback connectivity, related to alpha frequencies in occipital regions, and altered in neurological and psychiatric conditions ^8,9,15,17,19,20^. At the level of singletimecourses, we found that psilocybin altered contextual brain responses, specifically reducing surround suppression (Fig 2d, emphasised by black arrows). At the level of entire ROIs, we found that psilocybin significantly reduced surround suppression in early visual field maps (V1-3), while leaving activations largely unchanged (Fig 2e-g, emphasised by black arrows. Time points with black outline indicate *p*<0.01 difference with respect to placebo). Representative responses for all visual field maps and clusters are shown in Extnded Data 3. In sum, we found that psilocybin altered contextual cortical responses to visual stimuli, specifically reducing surround suppression in early visual field maps. This finding highlights the alteration of contextual computations as a potential mechanism underlying the effects of psilocybin in the human brain, and demonstrates a novel neuromodulatory contribution to surround suppression.

**Fig. 2.**
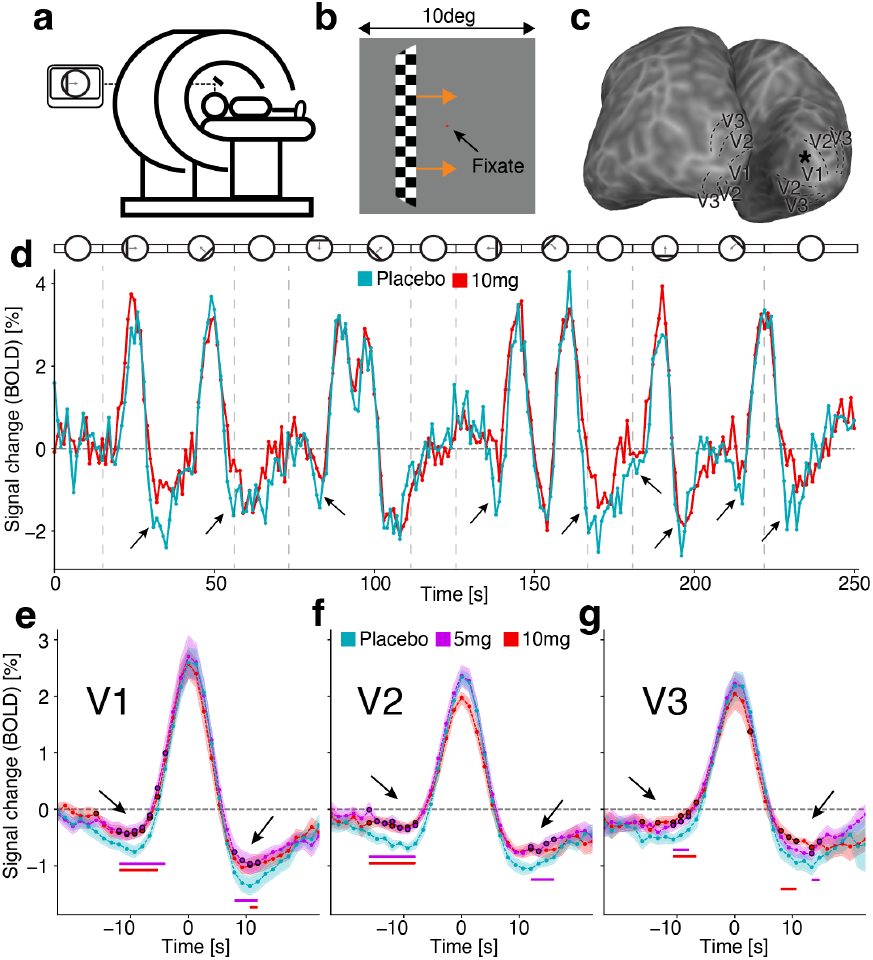
Psilocybin alters contextual brain responses, reducing surround suppression in early visual ROIs. **a**, 7T fMRI responses were recorded for all participants while viewing a simple visual stimulus (moving bar) in individual sessions with placebo, 5mg, and 10mg psilocybin doses in randomized order. **b**, Participants were instructed to fixate and performed a dot-color change task at fixation while a checkerboard bar moved through the screen in TR-locked steps in 8 different directions across the central 10 degrees of visual field and eye movements were recorded (Extended Data 2). **c**, Responses were resampled to individual anatomical surfaces obtained, and visual ROIs defined on the basis of individual polar angle and eccentricity maps. Asterisk on cortical surface indicates location of the timecourse shown in **d**, Example timecourse from a single participant, single cortical location in V1, in placebo and 10mg psilocybin doses. On top, stylized representation of moving-bar stimulus. At the single timecourse level, response activation was near-identical in placebo and 10mg psilocybin doses. surround suppression, the negative deflections of timecourse flanking the central positive activation peak, was systematically reduced in 10mg doses (highlighted by arrows). At the visual-field-map level, **e**, Representative responses in V1 (centered with respect to each bar-pass and averaged) **f**, V2 and **g**, V3 showed significant reduction of surround suppression while activation was not significantly altered. Timepoints with statistically significant differences (*p*<0.01) with respect to placebo are shown as circles with a black outline (Fisher permutation test, two-sided, 10^6^ permutations). Consecutive timepoints with significant differences are underlined with the color of the respective dose condition.

## Computational model captures psilocybin effects

To model cortical responses and the changes induced by psilocybin, we used a recently introduced population receptive fields (pRF) model ^9,14^ based on divisive normalization (Fig 3a, Eq 1). We have previously shown that the model captures a variety of contextual modulations observed in cortical responses such as surround suppression and nonlinear spatial summation thanks to local variation in its modulatory parameters, activation and normalization constants (Fig 3a,b Eq 1 parameters b and d) ^15^. In particular, the activation constant (parameter b in Fig 3a,b and Eq 1) modulates surround suppression (Fig 3b). Identically to ^15,21^, we fit model parameters to maximize variance explained at each cortical location, for each participant, in placebo, 5mg, and 10mg doses (Methods). The changes elicited by psilocybin in cortical responses were particularly evident in the reduction of surround suppression in early visual field maps, while activations were near-identical (Fig 2e-g). Given this pattern of results, we expected estimates of the activation constant should be reduced in psilocybin conditions relative to placebo. Confirming this hypothesis, we found individual cortical maps of model variance explained and visual-field eccentricity remained near-identical under psilocybin (Fig 3c-h). We found no significant alterations in model variance explained, activation pRF size, normalization pRF size, and normalization constant in V1 (Fig 3i-l). We found a significant, systematic decrease in the model activation constant in early visual field maps (V1-3) in both psilocybin doses (Fig 2m). This result could not be explained by potential concurrent changes in noise or hemodynamic response shape, which was also fitted at each cortical location, and verified with a crossvalidated approach (Extended data 4). The full model parameter estimates for all visual field maps and clusters are reported in (Extended data 5). In sum, a model-based approach provided additional evidence that psilocybin altered contextual computations in cortical responses, particularly in the form of reduced surround suppression. The model algorithmically explained psilocybin effects through variation in a specific modulatory parameter (the activation constant, parameter b in Fig 3a,b and Eq 1).

**Fig. 3.**
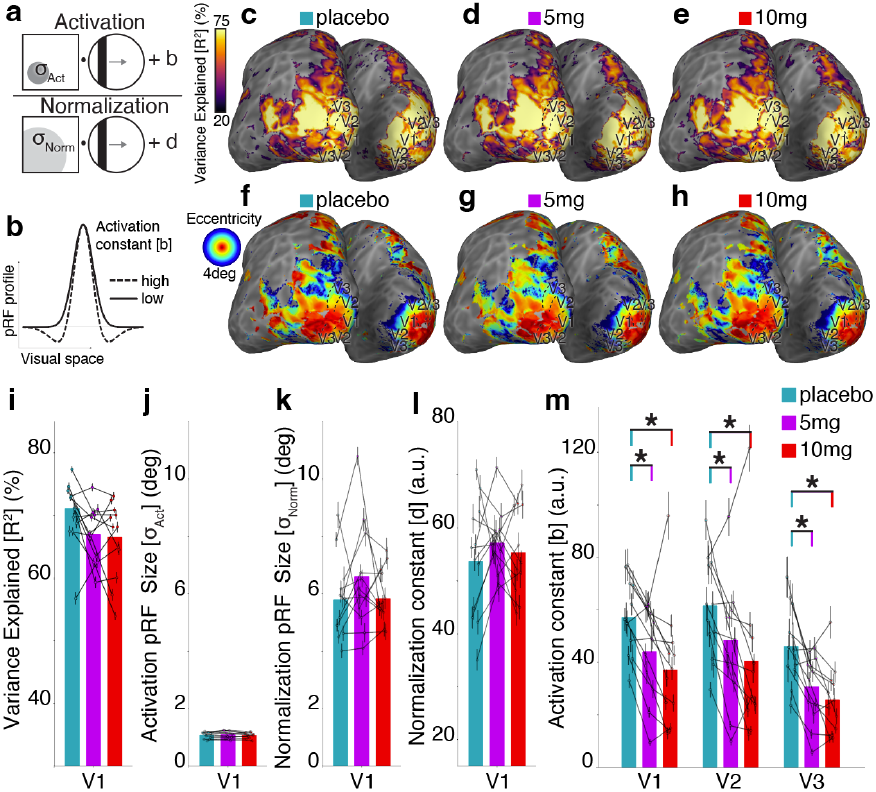
Computational model captures psilocybin effects on contextual brain responses through variation in specific modulatory parameter. **a**, Schematic of the DN pRF model (Eq. 1). The model prediction is defined as the ratio of activation and normalization components, convolved with a fitted hemodynamic response function (Methods). Activation and normalization terms are obtained as the dot-products of 2D Gaussians in visual space with the stimulus. At each cortical location, the optimal model prediction is obtained by maximizing Variance Explained (*R*^2^) as a function of the model parameters. **b**, The activation constant (parameter b) modulates surround suppression. A higher value of the activation constants (dashed profile) increases surround suppression. **c**, Example individual cortical maps of variance explained in placebo, **d**, 5mg, and **e**, 10mg showing the model’s ability to capture variance in brain responses is near-identical. **f**, Example individual cortical maps of eccentricity in placebo, **g**, 5mg, and **h**, 10mg showing the basic retinotopic structure of brain responses is near-identical. Psilocybin did not significantly alter **i**, variance explained **j**, activation pRF sizes, **k**, normalization pRF sizes, nor **l**, the normalization constant in V1. **m**, Psilocybin significantly and systematically decreased estimates of the activation constant (parameter b) in early visual field maps (V1-3), consistent with the observation of reduced surround suppression. Asterisks indicate *p*<0.01 (Fisher permutation test, two-sided, 10^6^ permutations).

Psilocybin, like other classic psychedelics, causes a variety of dose-dependent visual subjective phenomena ^22^. We hypothesised that the observed reduction in surround suppression might be a key mechanism underlying the subjective visual phenomena elicited by psychedelics. In this “unsuppressed” state, the visual system would allow increasing amounts of activity, normally quenched by surround suppression, to instead persist and propagate. Consistent with this hypothesis, we found significant correlations between the psilocybin-induced change in V1 estimates of the model activation constant, and subjective intensity of visual phenomena elicited by psilocybin (Extended Data 6). In particular, the change in activation constant significantly correlated with ratings of questionnaire items related to classic psychedelic visual phenomena, such as ‘I see geometric patterns’ and ‘I see movement in things that aren’t really moving’ (Extended Data 6), while generally lacking correlation with non-visual questionnaire items (Extended Data 8). We hypothesise that the change in the model activation constant represents an algorithmic hallmark of a more general effect of psychedelics on contextual computations, and particularly on surround suppression, manifested here in early visual field maps.

Surround suppression is a particular type of contextual modulation, involving both circuitry and neuromodulatory components ^7,8^. A key function of surround suppression is to increase tuning precision by contextually decorrelating responses ^8,23^. Previous studies have found that the anatomical size of V1 (inversely) correlates with the Ebbinghaus illusion ^12,24^. The increased degree of response correlation and functional overlap implied by a smaller cortical space being allocated to the same visual field have been suggested to be the relevant underlying functional properties ^11,13,25^. Here, we found that the Ebbinghaus illusion is increased by psilocybin (Fig. 1), while surround suppression is reduced in cortical responses (Fig. 2), an effect captured by a decrease in the activation constant of the normalization model (Fig. 3). The observed reduction in surround suppression implies an overall increase in functional response overlap, and reduction in tuning precision. As such, we hypothesised that the psilocybin-induced change in surround suppression could underlie the change in Ebbinghaus illusion we observed here. Consistent with this hypothesis, we found significant correlations between the psilocybin-induced change in V1 estimates of the model activation constant, and the increase in the Ebbinghaus illusion (Extended data 6). A well-known characteristic of psychedelic states is an increased sensitivity to the current context ^26^. We hypothesise that the altered contextual modulation in the Ebbinghaus illusion represents a perceptual hallmark of a more effect of psychedelics on contextual computations, manifested here in the visual domain.

We chose to use a computational model of population receptive fields based on divisive normalization ^7,15^. The changes we observed in brain responses appeared primarily in the form of surround suppression, and primarily in early visual ROIs (Fig. 2). Other models capable of capturing surround suppression, such as the Difference of Gaussians model ^17^, could have also been used. However, the normalization model is uniquely able to simultaneously capture a variety of contextual modulations, not limited to surround suppression ^15^. The normalization model outperforms other models throughout the human visual system, including the DoG model in early visual ROIs ^15^. The modulatory parameters of the normalization model correlate with neurotransmitter receptor densities, demonstrating ties with the underlying biology ^21^. Other contextual modulations, such as nonlinear spatial summation, could have potentially been concurrently affected by psilocybin. If so, they would have confounded models unable to capture nonlinear contextual modulations, such as the DoG model. The normalization model allowed us to ensure that the changes we observed in the data were specific to surround suppression, and not confounded by additional potential changes in other contextual modulations. For these reasons, we concluded the normalization model would be the best-available choice to capture the changes induced by psilocybin.

Visual spatial computations provide a general scaffolding for cognition ^27^, with the early visual cortex acting as a “multiscale cognitive blackboard” ^28^. Visual spatial population receptive fields are present in occipital, parietal, ventral, temporal, frontal areas ^9,16,29,30^, and have recently been observed even in regions generally thought to be amodal, such as the default mode network ^31^, at the interface of perception and memory processing ^32^, in the cerebellum ^33^, and in the hippocampus ^34^. Alterations in lower levels of the brain’s foundational visual-spatial cognitive architecture, such as those we observe here, are hence likely to have far-reaching implications on brain dynamics, perception, and subjective experience.

Alterations in contextual computations appear as an underlying thread linking our findings to existing clinical, phenomenological, and neuroscientific lines of evidence. Contextual computations ^35^, divisive normalization ^7^, surround suppression ^8^, and receptive fields ^6^ are ubiquitous and fundamental properties of cortical responses in a variety of sensory and cognitive domains. Contextual factors have long been known to play a crucial role in psychedelic experience and therapeutics ^5,26^. Studies in animal models have shown that psychedelics reopen a critical period for contextual reward learning ^36^. Psychedelics show promise in the treatment of depression ^37^. Depression has been characterized as stemming from an impaired ability to combine current input with contextual factors ^35^. In light of the existing literature, our findings highlight the alteration of contextual computations as a potential general computational mechanism underlying the action of psychedelics in the human brain. We speculate that the compound effects of an apparently simple computational alteration, ubiquitously present throughout sensory and cognitive domains and at various stages of the brain’s functional hierarchy, might hold the potential to underlie the diversity and apparent paradoxicality of psychedelic effects.

Our study is limited in several respects. First, the BOLD signal is determined by both neuronal and hemodynamic factors, which may also be altered by psilocybin. To address this directly, we fitted the HRF shape to vary at each cortical location, for each participant, at each dose, ensuring that hemodynamic changes are captured separately from computational changes. We also carried out a crossvalidated comparison of models including purely neural changes, purely hemodynamic changes, both, and neither. These analyses showed that the alterations in cortical responses elicited by psilocybin could not be explained by potential concurrent changes in noise or hemodynamics (Extended Data 4,5). Second, we investigated 5mg and 10mg doses of psilocybin, while previous neuroimaging studies in healthy and clinical populations have generally investigated higher doses (up to 25mg) ^38^. At higher doses, it is likely that impairments in task performance or fixation ability would confound the assessment of computational changes elicited by psilocybin. Our results showed that intermediate doses of psilocybin are sufficient to induce systematic changes in perception and cortical responses, without significantly impairing fixation ability or performance on simple tasks (Extended Data 2). Third, psilocybin and its active metabolite psilocin bind to multiple receptors, including both 5-HT2A and 5-HT1A. Our study was not designed to adjudicate if an individual receptor (or a combination thereof) was responsible for the effects of psilocybin we observed. Nonetheless, this question may be addressed using our approach in combination with chemical blockers of individual receptors. Fourth, we focused on spatial vision, allowing psilocybin’s effects on visual-contextual computations to unequivocally manifest. This approach could be extended in order to assess the alteration of contextual computations as a potential general computational mechanism in other sensory and cognitive domains.

In this study, we found that psilocybin alters contextual perception in the Ebbinghaus illusion, contextual modulations in brain responses to visual stimuli, and proposed a computational model capable of capturing and linking these changes. We concluded that psilocybin alters visual-contextual computations. Our findings highlight the alteration of contextual computations as a potential general and parsimonious mechanism underlying the effects of psychedelics on brain and behavior.

## Materials and Methods

### Recruitment and exclusion criteria

Participation was solicited via public online means. Interested participants completed an online form, which was used as an initial screening for exclusion criteria. Participants received a small monetary compensation for their participation in the study. Exclusion criteria: age below 21 years, age above 55 years, no prior experience with hallucinogens, previous adverse reactions to hallucinogens, previous adverse responses to MRI, personal or family history of psychiatric or neurological conditions, ongoing use of medications or recreational drugs. Participants were asked to refrain from the use of recreational drugs in the four weeks leading up to the experiment and between the experimental session. Participants that did not meet any exclusion criteria based on the online questionnaire responses were invited to the Spinoza Centre for further in-person screening. Participants who did not meet any exclusion criteria based on online and in-person screening participated in a preliminary data collection session. During this session, we collected two anatomical scans, one functional scan, and ensured participants had normal fixation ability and eye movement traces and normal or corrected-to-normal visual acuity. Two prospective participants were excluded at this screening stage, before being enrolled in the study, because of the presence of potentially abnormal eye movements (nystagmus). This selection process ensured that all participants were not naive to MRI scanning and to the experimental tasks, prior to confirming their enrollment in the study.

### Participants

Eighteen participants (aged 22 to 45 years, eight female) were enrolled in the study. All participants had normal or corrected-to-normal visual acuity. All studies were performed with the informed written consent of the participants and were approved by the Medical Ethics Committee (METC) of the Amsterdam University Medical Centre. Data from five participants were excluded from analysis because of the presence of scanner artifacts (coil failures) during one or more of their sessions. Data from one participant were excluded from the analysis due to excessive head motion in one session. A total of twelve participants were included in the final analysis of fMRI data.

### Experimental design

The experiment followed a randomized, double-blind, placebocontrolled, crossover design. Each participant underwent three experimental sessions and was administered placebo, 5mg, and 10mg of psilocybin, in random order, at least two weeks apart from each other.

### Psychophysical model

In the Ebbinghaus illusion, participants were asked to fixate on a central cross and presented white circles on an isoluminance background for a short interval (0.4s). For each presentation, they were asked to report via button press whether they perceived the right or left circle as larger. In control trials, the two circles were presented in isolation. In test trials, one of the circles was presented surrounded by a set of larger circles. We estimated a standard psychophysical model, in which the perception of the two stimulus features (the radius of the reference stimulus *r*_*r*_ and the embedded stimulus *r*_*e*_) are formally described as samples from two normal distributions:

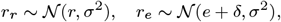

where *r* is the actual radius of the reference stimulus and *e* is the actual radius of the embedded stimulus; *σ*^2^ quantifies the inherent perceptual noise due to sensory processing limitations or neural variability in representing the stimulus radii; *δ* captures systematic biases in perceiving the radius of the embedded stimulus, potentially driven by contextual effects in the Ebbinghaus illusion. The probability that a participant indicates the reference stimulus is larger than the embedded stimulus can be described by their difference distribution:

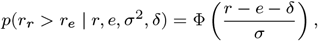

where Φ is the standard normal cumulative density function.

To account for lapses, we also estimated the proportion of trials *p*, on which the participant did not process the stimulus and responded randomly:

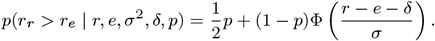

The noise parameter *σ*^2^, the bias parameter *δ*, and the lapse parameter *p* were estimated in a hierarchical framework using the No U-Turn (NUTS) Hamiltonian MCMC sampler as implemented in pymc ^39^. The psychometric model was implemented in the Python package bauer ^40^.

For a given parameter *θ* (e.g., *σ*^2^ or *δ*), the participant-session-specific parameter of participant *p* at session *t* was modeled as:

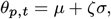

where *µ* is the group mean parameter, *σ* is the standard deviation of the group distribution, and *ζ* is an individually determined offset parameter. This offset-based specification of the hierarchical model was chosen to prevent likelihood “funnels” that affect high-dimensional hierarchical models ^41^.

To estimate the parameters separately for the three dose conditions (placebo / 5mg / 10mg), with and without surround, we used a regression approach ^42^. A given parameter *θ*_*p,s,g*_ (e.g., *δ* or *σ*^2^) for observer *p* on a trial *t* and dosage *d* was modeled as:

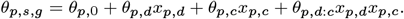

Where *x*_*p,c*_ indicated the condition (with or without surrounding circles) of a trial and *x*_*p,d*_ the dosage (0/5/10mg). The values of (and the uncertainty in) the estimates of *θ* could then be used to quantify the relative effect of the different doses, with and without surround. We used mildly informative priors on all group distribution parameters: For the precision parameter *σ*:

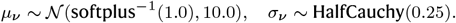

Because *σ* should be bounded between [0, ∞), we estimate it via *ν* on the unbounded scale and then brought it to the bounded scale using the softplus function to evaluate the likelihood function: *σ* = softplus(*ν*), where softplus(*x*) = ln(1 + *e*^*x*^) is the softplus function and softplus^−1^ is its inverse.)

For the bias parameter *δ* we used the following hierarchical parameters

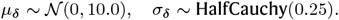

For the lapse parameter *p*:

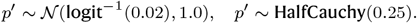

(Here, *p* was bounded between 0 and 1 by using the logit function: 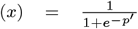, and logit^−1^ is the logistic function. The median of our prior on *p* was thus 0.02.)

### Anatomical MRI scans

T1-weighted and T2-weighted structural MRI data were acquired using a Philips Achieva 7T scanner with an 32-channel Nova Medical head coil, at a resolution of 0.7 mm isotropic. Freesurfer 7.2 recon-all was used to obtain native cortical surface reconstructions for each participant ^43^. The software makes use of the T2w image to refine the segmentation obtained by T1w alone, particularly in the exclusion of the sagittal sinus and at the pial surface border. Anatomical segmentations were further refined manually using Freesurfer’s software Freeview ^43^.

### Functional MRI scans

Functional MRI data were acquired using a Philips Achieva 7T scanner with an 32-channel Nova Medical head coil. Functional scans were carried out with a 3D EPI sequence, with a repetition time (TR) of 1.32s, and spatial resolution of 1.79mm isotropic. The first 10s of recorded data at the beginning of each scan were automatically discarded by the scanner to avoid start-up magnetization transients. Each pRF mapping scan lasted for a total of 255 TRs (approximately 336s). For each participant, on each experimental session, we collected six pRF mapping scans. At the end of each scan, a top-up scan with opposing phase-encoding direction was recorded, to perform susceptibility distortion correction.

### Functional MRI data preprocessing

Raw scanner data was converted to nifti using dcm2niix and stored in BIDS format ^44^. Thermal denoising was applied using the NORDIC algorithm ^45^. Subsequent functional preprocessing steps (susceptibility distortion correction, coregistration, resampling of volumetric data to fsnative surfaces) were performed using fMRIprep v20.7 ^46,47^. The first 20 principal components of the anatomical and temporal physiological regressors computed by fMRIprep were regressed out using pybest ^48^. Finally, BOLD timecourses in fsnative surface spaces were converted to percent signal change and highpass filtered to remove drifts using a cosine filter (regressing out the first three components).

### Head motion

We estimated motion parameters using MRIQC ^49^. We defined excessive motion as a mean frame displacement across scans in a session exceeding one third of the voxel size (0.6mm) or a mean of the maximal frame displacement across scans in a session exceeding one voxel size (1.8mm).

### Population receptive field modeling

Population receptive field modeling followed the same procedure as ^15,21^ and was carried out using dedicated software prfpy ^50^ and prfpytools ^51^. We used a standard pRF mapping stimulus, a drifting checkerboard bar ^14,15^. The stimulus aperture subtended 10deg of visual angle. The checkerboard pattern inside the bar moved parallel to the bar orientation, and the bar itself stepped in the perpendicular direction at every TR (1TR=1.32s). Eight bar configurations were presented (two cardinal and two diagonal directions, in two motion directions). The width of the bar subtended 1.25deg of visual angle; the bar swept across the stimulus aperture in 20 TR-locked equal-size steps. A period of 15 TRs of mean luminance (0% contrast) was presented every two bar passes. Mean luminance periods of 15 TRs were presented at the beginning and end of each scan. A small fixation dot (0.1deg) was presented in the middle of the screen. This fixation dot changed color (red to green) at semirandom time intervals, and participants reported this change via button-press. The DN model equation was defined as ^15^

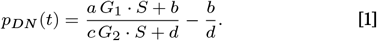

The spatial dependence of Gaussian pRFs *G* ≡ *G*(*x, y*) and the spatiotemporal dependence of stimuli *S* ≡ *S*(*x, y, t*) are omitted for brevity, and we denote:

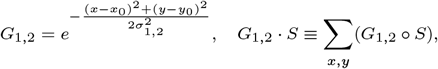

(*x*_0_,*y*_0_) is the response central location in the visual field, *σ*_1_, *σ*_2_ are the sizes of activation and normalization pRFs, *a, c* their amplitudes, *b, d* the activation and normalization constants. The neural prediction of the DN model *p*_*DN*_ (*t*) is then convolved with a hemodynamic response function to obtain a prediction of the BOLD fMRI signal at each cortical location. Identically to ^15,21^, the fitting procedure starts by fitting a Gaussian pRF model, with a grid-search stage followed by an iterative stage, to obtain initial estimates of pRF size and position. These estimates are used as starting parameters for the DN model fit, which also comprises of a grid-search and iterative fit stages. First, a grid-search is performed vertex-wise for the additional DN model parameters, while keeping the pRF size and position fixed to the Gaussian estimates; next, an iterative fit over all the DN model parameters is performed. Both grid-search and iterative fit stages of both models include a one-parameter hemodynamic response function, computed as a linear combination of the standard SPM-software HRF and of its derivative; the fit parameter represents the coefficient of the HRF derivative.

### Visual regions of interest and clusters

Visual ROIs and clusters were defined for each participant on the basis of individual polar angle and eccentricity maps of cortical responses, following ^16,52^, and similarly to ^15^.

### Population receptive field parameters estimates

Parameter estimates for each ROI were obtained as the *R*^2^-weighted mean of parameter estimates at each cortical location within the ROI. To prevent edge artifacts or low-signal timecourses from affecting estimates, cortical locations with model *R*^2^ *<* 0.3 or eccentricity outside the bounds of the stimulus range were excluded.

### Representative ROI timecourses

Representative ROI timecourses were obtained as the *R*^2^-weighted mean of timecourses at each cortical location within the ROI, split for each bar-pass, temporally aligned with respect to the pRF position (*x*_0_,*y*_0_) and accounting for HRF delay. To prevent edge artifacts or low-signal timecourses from affecting estimates, cortical locations with model *R*^2^ *<* 0.5, difference between DN and Gaussian *R*^2^ *<* 0.1, or eccentricity outside the bounds of the stimulus range were excluded.

## Acknowledgements

The authors would like to acknowledge the experiment participants for their dedication and support to the study. MA would like to acknowledge Jurjen Heij for support with preprocessing pipelines, Tom Pelletreau-Duris for support in data analysis, and Leon Behle for support in stimulus design and testing.

## Dataand Materials Availability

All data needed to evaluate the conclusions in the paper are present in the paper and/or the Supplementary Materials.

## Author Contributions

Conceptualization: MA, TK, SD. Methodology: MA, GH, NV, TK, SD. Investigation: MA, GH, NV, TK, SD. Visualization: MA, GH, NV. Supervision: TK, SD. Writing, original draft: MA. Writing, review & editing: MA, GH, NV, TK, SD.

## Additional information

### Supplementary information

Provided in dedicated submission file.

