## Extended Data 1 for "Psilocybin alters visual contextual computations"

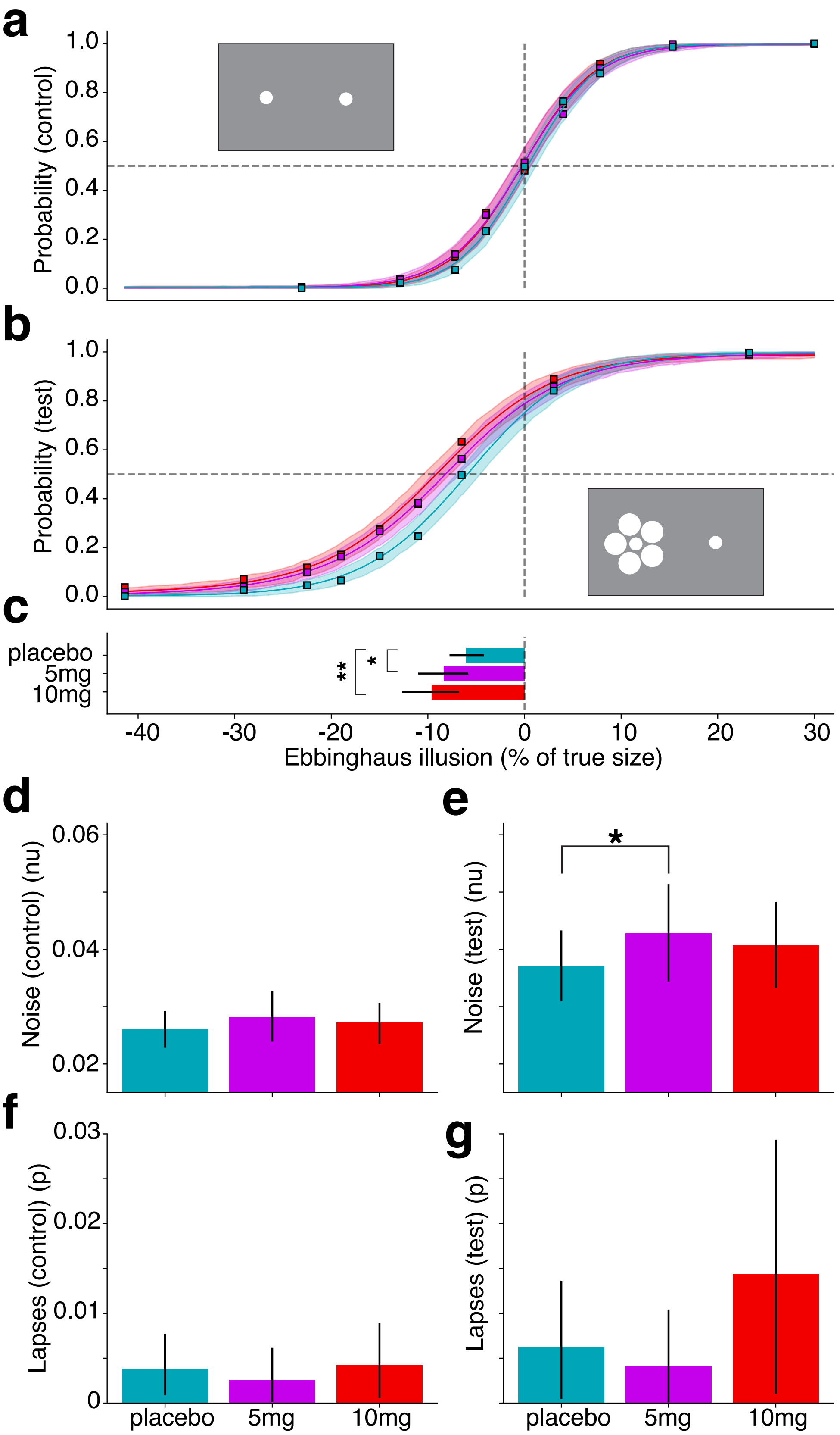


**Psychophysical data and model parameters for Ebbinghaus illusion and control conditions. a,** Psychometric curves in control condition (i.e. size-judgement without the presence of contextual stimuli). **b,** Psychometric curves in test condition (i.e. size-judgement with the presence of contextual stimuli) showing the Ebbinghaus illusion. **c,** Bias parameter estimate for placebo, 5mg, and 10mg doses in test condition (5mg: p=0.012, 10mg: p=0.005) showing psilocybin significantly increases the Ebbinghaus illusion. **d,** Noise parameter (nu) estimate for placebo, 5mg, and 10mg doses in control condition showing no significant differences relative to placebo (5mg: p=0.13, 10mg: p=0.21). **e,** Noise parameter (nu) estimate for placebo, 5mg, and 10mg doses in test condition. The noise parameter was increased with mild significance in 5mg dose (5mg: p=0.044) but not in 10mg dose (10mg: p=0.11). **f,** Lapses parameter estimate for placebo, 5mg, and 10mg doses in control condition showing no significant differences relative to placebo (5mg: p=0.76, 10mg: p=0.47). **g,** Lapses parameter estimate for placebo, 5mg, and 10mg doses in test condition showing no significant differences relative to placebo (5mg: p=0.72, 10mg: p=0.13). In sum, psilocybin signifcantly increased the Ebbinghaus illusion both in 5mg and 10mg conditions. The only other significant effect observed was an increase in noise in the test condition for the 5mg dose. The psychophysical model (Methods) allowed us to distinguish the effect of psilocybin on the Ebbinghaus illusion from any concurrent effects on noise or lapses.
