## Extended Data 2 for "Psilocybin alters visual contextual computations"

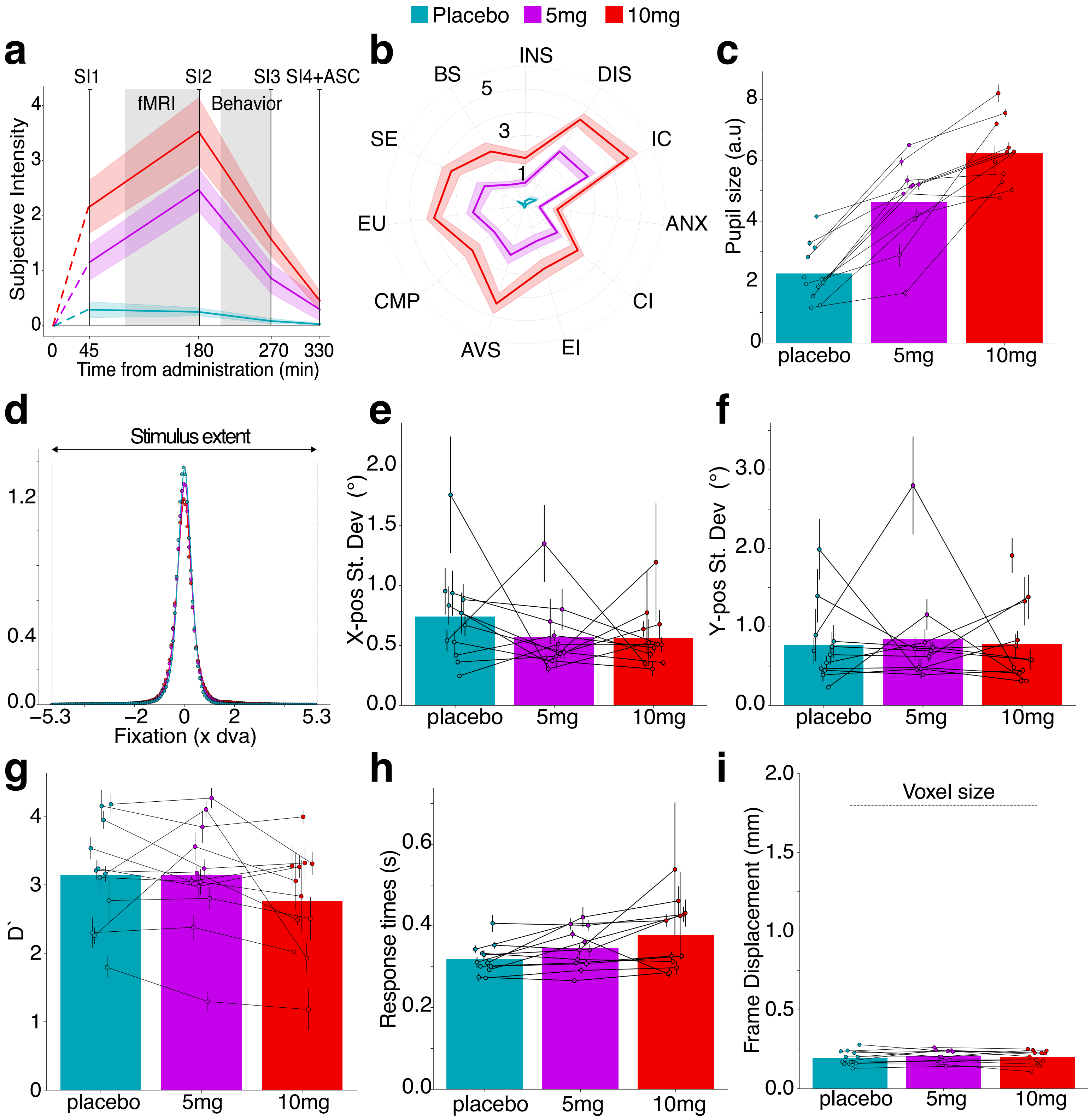


**Subjective intensity, pupil size, fixation stability, task performance, and head motion during MRI scans with placebo, 5mg, and 10mg psilocybin.** Unless otherwise stated, all statistical testing for these data was conducted using ANOVA. **a,** Timecourse of subjective intensity assessed with 21-question subjective experience questionnaire at four timepoints. The effect of psilocybin was statistically significant at 45min, 180min, and 270min post administration ($p < 10^{-2}$). At the last timepoint, approximately 330min post administration, subjective experience was statistically indistinguishable from placebo ($p = 0.27$). **b,** Retrospective assessment of subjective intensity with 11-dimension Altered States of Consciousness questionnaire. The effect of psilocybin dose on 11D ASC was statistically significant in all dimensions (all $p < 10^{-2}$), with the exception of Disembodiment and Blissful State, which were near-significant ($p = 0.014$ and $p = 0.012$, respectively). **c,** During pRF-mapping fMRI scans, we collected pupil-size measurements and gaze position using an EyeLink 1000 (SR Research Ltd.). We found a statistically significant increase in pupil size ($p < 10^{-2}$). **d,** Horizontal-axis fixation density, relative to stimulus extent, showed qualitatively good fixation in all conditions. Quantitatively, **e,** Horizontal-axis standard deviation of fixation during MRI scans and **f,** Vertical-axis standard deviation of fixation were not significantly altered by psilocybin ($p = 0.49$ and $p = 0.75$, respectively). We conclude that pupil size, but not fixation ability, was significantly altered by psilocybin at 5mg and 10mg doses. **g,** During pRF-mapping fMRI scans, participants performed a simple dot-color change task at fixation, reporting via button-press. We observed no statistically significant differences in participants performance, quantitatively indexed by D′ ($p = 0.06$). **h,** The temporal distribution of responses, quantitatively measured by the distribution mode, was slightly slower with psilocybin, near significance (p = 0.013). We conclude that participants were not only able to fixate (panels d-f) but also to complete a fixation color-change task in all three conditions with satisfactory performance, up to a trend-level delay in response times. **i,** Frame displacement (head motion) during MRI scans was not significantly altered by psilocybin (p=0.45).
