## Extended Data 3 for "Psilocybin alters visual contextual computations"

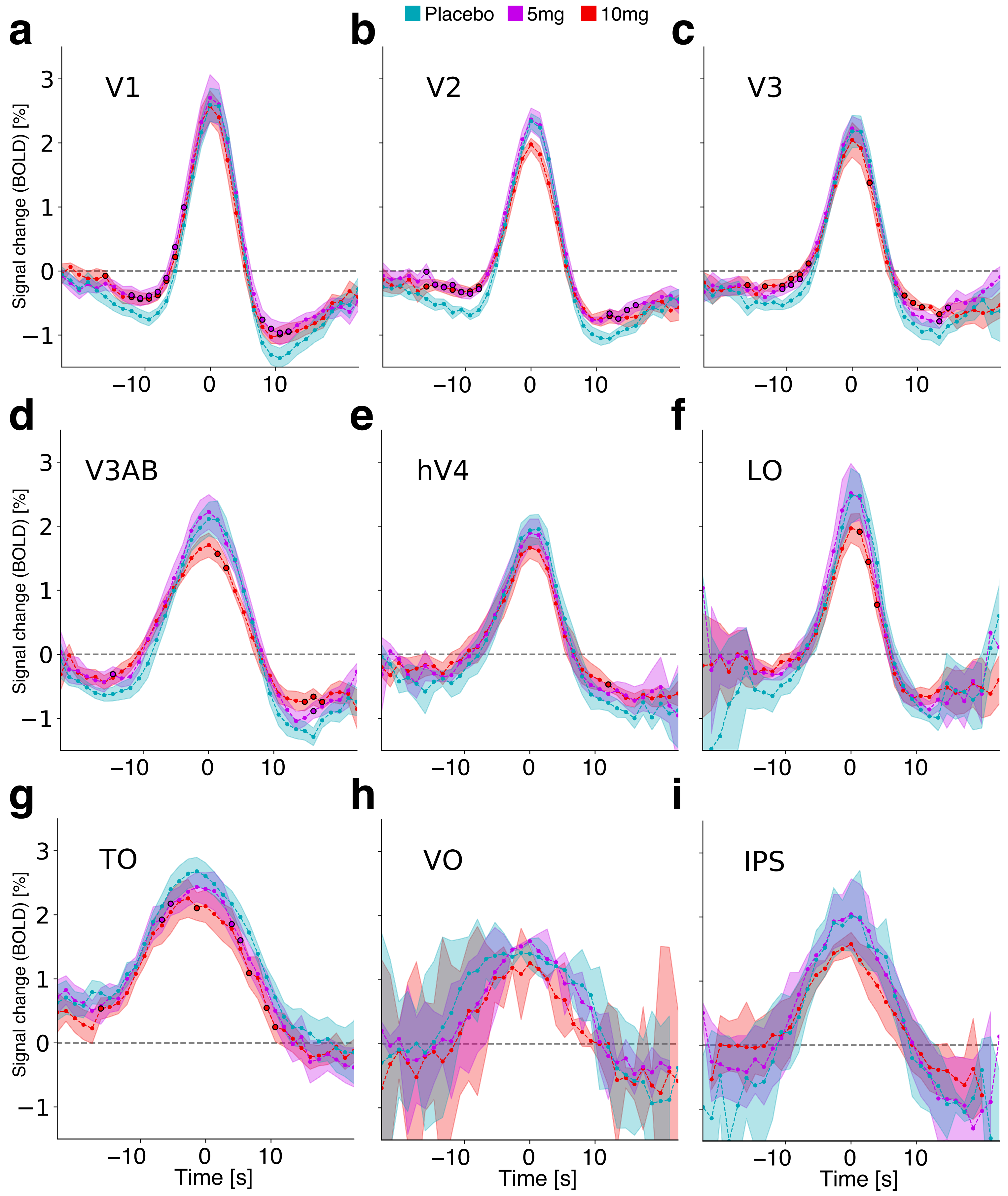


**Bar-pass-aligned, averaged response for visual ROIs and clusters.** Visual ROIs and clusters were defined for each participant on the basis of individual polar angle and eccentricity maps. Responses to individual bar-passes at each vertex were aligned relative to pRF positions, and averaged across ROIs and participants to obtain representative responses for each visual ROI and cluster. At each timepoint, statistical testing was conducted with Fisher permutation test ($10^6$ permutations). Timepoints with statistically significant differences ($p<10^{-2}$) with respect to placebo are indicated with a black circle outline. Early visual ROIs **a,** V1, **b,** V2, and **c,** V3 showed a near-identical pattern of results, with little or no significant differences in activation, but significant and systematic differences in surround suppression. Intermediate visual ROIs and clusters **d,** V3AB, **e,** hV4, and **f,** LO showed little to no systematic differences, up to a potential trend-level reduction in surround suppression, and a slightly reduced response amplitude/-faster decay in LO. Late visual ROIs and clusters **g,** TO, **h,** VO, and **i,** IPS showed little to no significant differences, up to a slightly reduced response amplitude/faster decay in TO, similar to the neighboring LO cluster. We conclude that psilocybin primarily acts in the visual system by exerting a systematic and significant reduction of surround suppression in early visual ROIs, up to a potential concurrent effect in late lateral and temporal occipital clusters.
