## Extended Data 4 for "Psilocybin alters visual contextual computations"

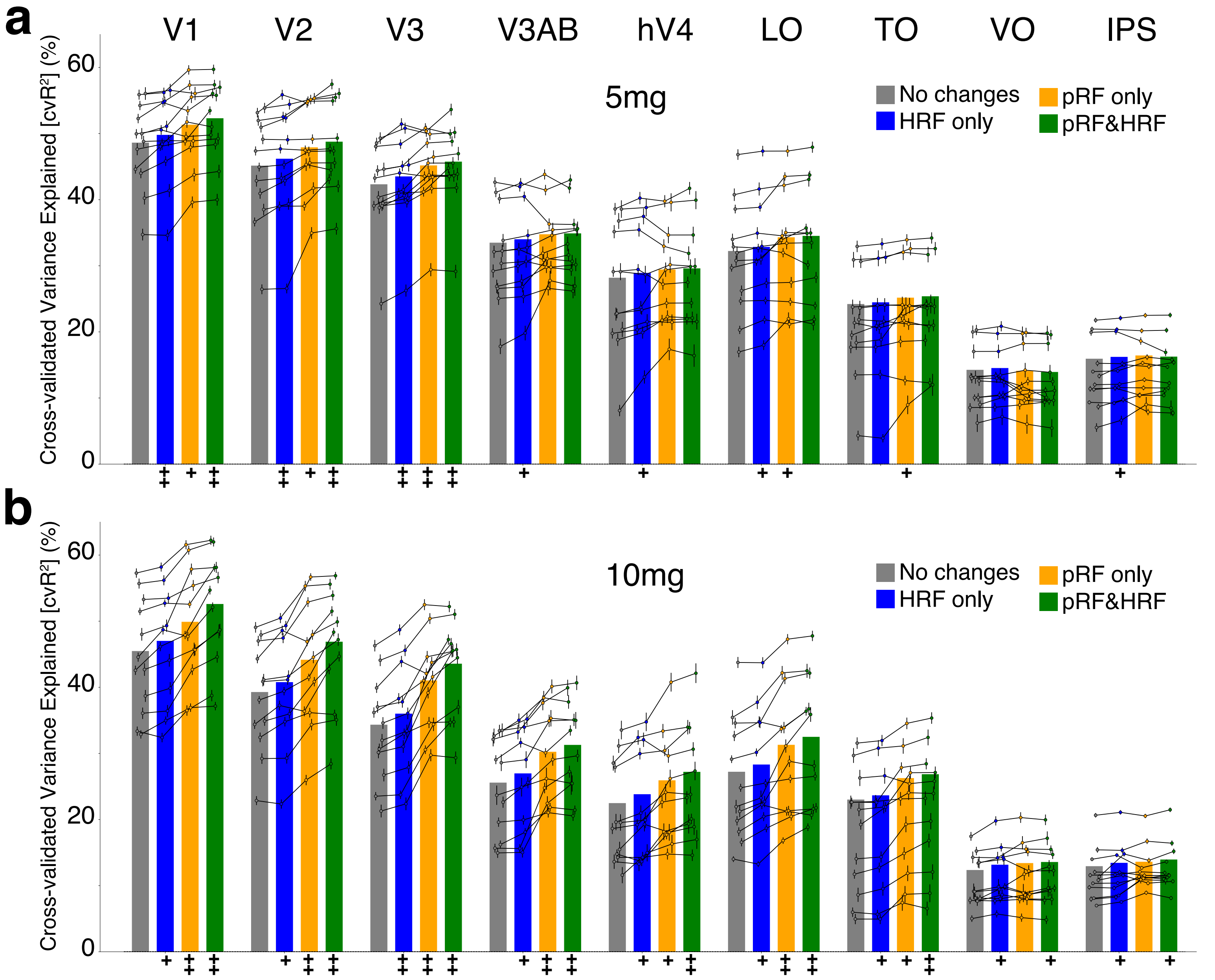


**Cross-validated analysis of variance explained by nested-model hypotheses.** We investigated whether the changes observed in data and computational model parameters in 5mg and 10mg psilocybin conditions might be explained by changes in noise or hemodynamics, rather than changes in signal. Signal, here, is intended as the amount of cross-validated variance explained that the computational model (DN-pRF) is able to capture. To do this, we compared four different models representing four different hypotheses of potential changes between placebo and each psilocybin dose. No changes (gray bars), only hemodynamic response changes (blue bars), only population receptive fields changes (yellow bars), both population receptive fields and hemodynamic response changes (green bars). We compared each hypothesis relative to the "baseline" (no changes) hypothesis using a Fisher permutation test ($10^6$ permutations). -/+:$p < 10^{-2}$, --/++:$p < 10^{-4}$, minus sign indicates significant decrease with respect to baseline, plus sign indicates significant increase. A statistically significant increase provides evidence that the relevant model hypothesis captures changes in signal not due to spurious factors. **a,** For the 5mg dose, significant differences are concentrated in early visual ROIs and confirm the presence of pRF parameter changes, consistent with the finding of reduced surround suppression and reduction in the DN model activation constant. The cross-validated analysis also indicates the presence of potential hemodynamic changes. In intermediate and late visual ROIs and clusters, the pattern of results suggest little to no changes, up to a potential hemodynamic change. **b,** For the 10mg dose, differences in early visual ROIs become larger in magnitude and significance. Significant differences can be observed also in intermediate visual ROIs and cluster, suggesting that at increasing doses psilocybin effects begin to "percolate" and affect areas further up the visual system hierarchy. Overall, this approach allowed us to disentangle the different aspects of psilocybin effects on population receptive field parameters and hemodynamic properies, confirming that the observed changes in pRF parameters are not explainable by noise or concurrent hemodynamic changes, as well as providing evidence that psilocybin exerts both neural and hemodynamic changes in the human brain. Future studies should consider including hemodynamic response modeling, as we do here, to disentangle these different effects.
