## Extended Data 5 for "Psilocybin alters visual contextual computations"

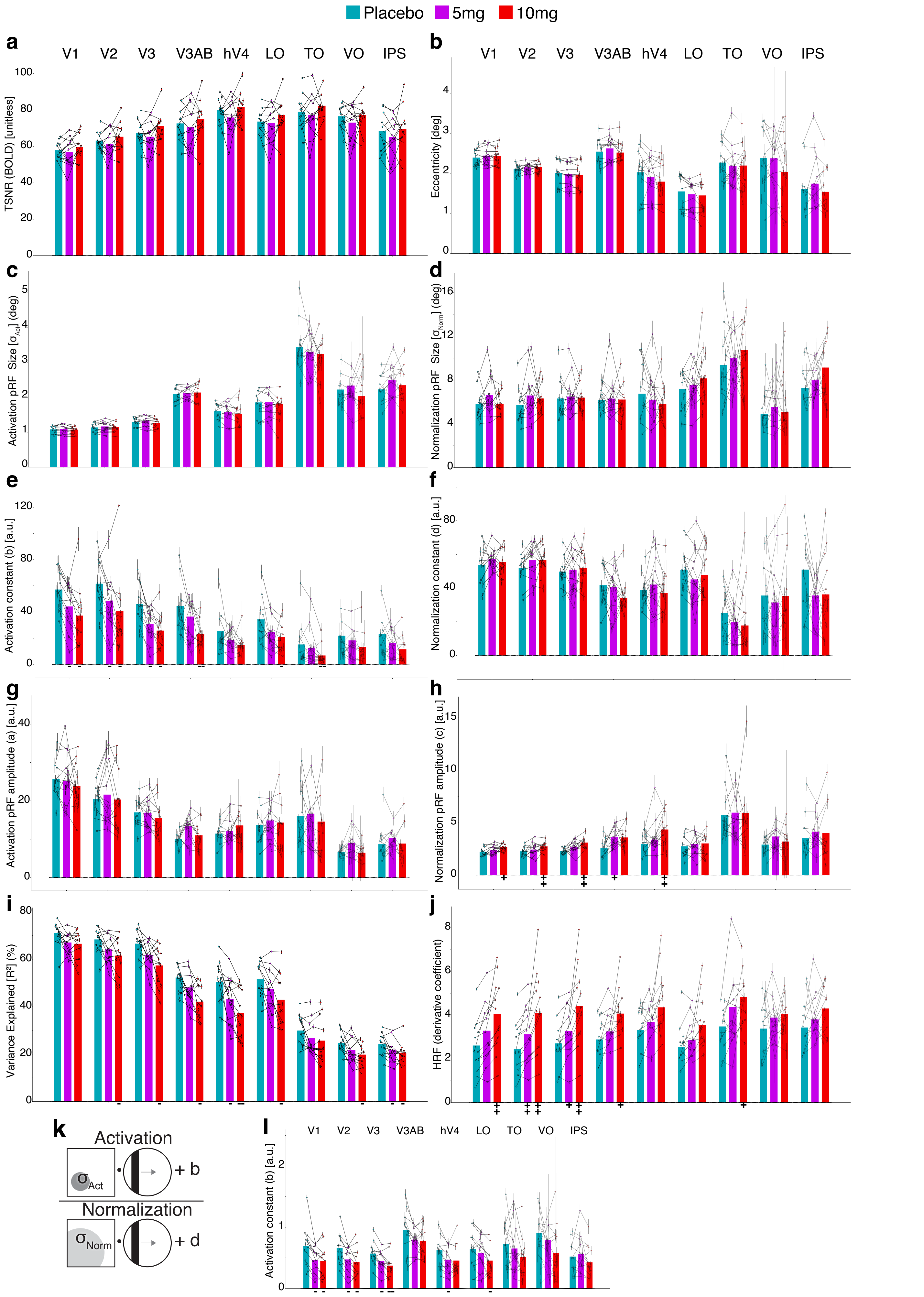


**DN model parameter estimates in visual system ROIs and clusters. a,** Signal to noise ratio i.e. signal standard deviation divided by the mean (TSNR). **b,** Eccentricity. **c,** Activation pRF size. **d,** Normalization pRF size. **e,** Activation constant. **f,** Normalization constant. **g,** Activation pRF amplitude. **h,** Normalization pRF amplitude. **i,** Variance expsplained. **j,** Hemodynamic function derivative coefficient. **k,** DN model equation. **l,** Activation constant estimated from alternative model formulation with d=1. For all panels, -/+:$p<10^{-2}$, --/++:$p<10^{-3}$, Fisher permutation test ($10^6$ permutations). Minus sign indicates significant decrease with respect to placebo, plus sign indicates significant increase.
