## Extended Data 6 for "Psilocybin alters visual contextual computations"

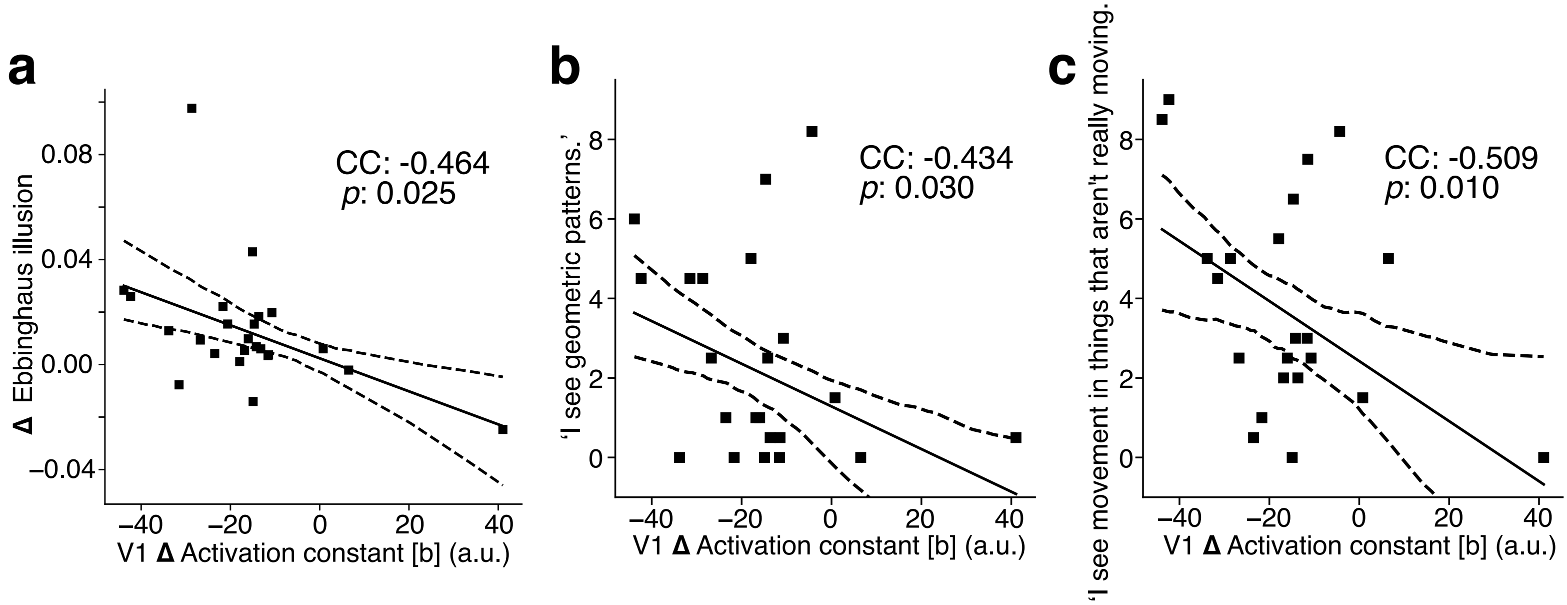


**Psilocybin-induced change in V1 model's activation constant correlates with change in Ebbinghaus illusion and subjective ratings of visual phenomena. a,** We hypothesised that the psilocybin-induced change in cortical surround suppression, captured by the DN model's activation constant, mediates the perceptual change induced by psilocybin in the Ebbinghaus illusion. Consistent with this hypohtesis, we found a correlation between the psilocybin-induced change in V1 activation constant, and the change in Ebbinghaus illusion. Previous studies have found that the anatomical size of V1 (inversely) correlates with the Ebbinghaus illusion. A smaller cortical space being allocated to the same visual field implies increased overlap between receptive fields, and consequent reduction in neural tuning precision. Surround suppression increases tuning precision by decorrelating responses and decreasing functional overlap between neighboring receptive fields. Hence, we hypothesised a reduction in surround suppression mediates the observed change in the Ebbinghaus illusion. **b,c,** We hypothesised that the psilocybin-induced change in cortical surround suppression, captured by the DN model's activation constant, might underlie subjective effects of psychedelics in the visual domain. Consistent with this hypothesis, we found a correlation between the psilocybin-induced change in V1 activation constant, and the subjective ratings of classic psychedelic visual phenomena such as 'I see geometric patterns.' and 'I see movement in things that aren't really moving'. Psilocybin, like other classic psychedelics, is well-known to cause a variety of dose-dependent visual phenomena. The reduction in surround suppression might be a key mechanism underlying some of the visual effects of psychedelics. In this unsuppressed state, the visual system is likely to allow increasing amounts of activity, normally quenched by surround suppression, to instead persist and propagate.
