## Extended Data 7 for "Psilocybin alters visual contextual computations"

**Subjective Experience assessment items**

To asses the temporal dynamics of subjective experience intensity, participants were asked to rate the following 21 items, in random order, on a 10-point scale, at 4 time points throughout each experimental day, adapted from previous studies (Pallavicini et al., 2021). The subjective experience intensity was then computed as the average rating of these items.

"I experience a disintegration of my 'self' or 'ego'."

"Things look strange."

"My experience has a spiritual or mystical quality."

"I feel a profound inner peace."

"I feel like I am floating."

"I experience a sense of merging with my surroundings."

"My thinking is muddled."

"My experience has a supernatural quality."

"I feel afraid."

"Edges appear warped."

"I feel unusual bodily sensations."

"I see geometric patterns."

"My sense of space and size is distorted."

"I feel suspicious and paranoid."

"I see movement in things that aren't really moving."

"My thoughts wander freely."

"My experience has a dream-like quality."

"My perception of time is distorted."

"My imagination is extremely vivid."

"Sounds influence what I see."

"I fear losing control of my mind."
