## Extended Data 8 for "Psilocybin alters visual contextual computations"

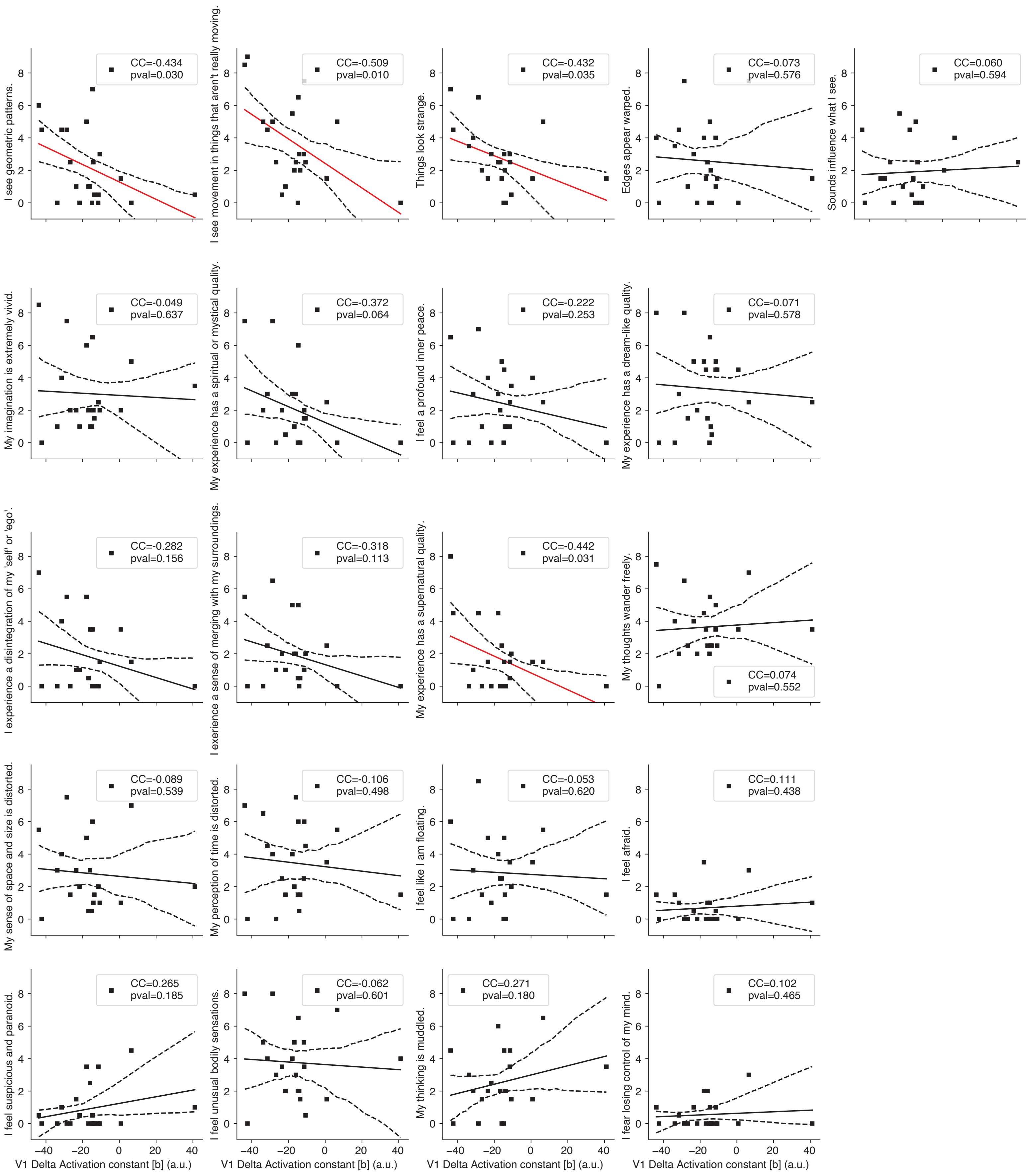


**Psilocybin-induced change in V1 model's activation constant correlates specifically with ratings of visual phenomena.** Correlation between psilocybin-induced change in V1 model's activation constant and ratings on all 21 items of subjective intensity questionnaire. The psilocybin-induced change in V1 model's activation constant does not broadly correlate with subjective intensity ratings, but particularly with visual items ('I see geometric patterns.', 'I see movement in things that aren't really moving', 'Things look strange.'). The only non-specifically-visual item with a significant correlation is 'My experience has a supernatural quality.'. (red fit-lines indicate $p<0.05$).
